# Classical music in cognitive rehabilitation: Three case studies of spatial neglect

**DOI:** 10.64898/2026.09.18.751660

**Authors:** Ryan Zeineldin, Yusaku Takamura, Maryane Chea, Lionel Naccache, Eléonore Bayen, Monica N. Toba, Paolo Bartolomeo

## Abstract

Chronic spatial neglect after right-hemisphere stroke reflects network dysfunction, with recovery influenced by structural connectivity. Music recruits coordinated bi-hemispheric activity and may promote interhemispheric communication after stroke. Following our pilot study in post-stroke aphasia, we asked whether classical music modulates connectivity and behavior in chronic neglect. Three patients completed a crossover protocol comparing two weeks of daily audiovisual classical music with an audiobook control, with neglect, mood, quality-of-life, diffusion MRI, and resting-state high-density EEG measures. All showed right superior longitudinal fasciculus II disconnection with preserved callosal pathways. In N2, line bisection shifted leftward across all five lines after music, together with increased posterior interhemispheric theta-band weighted symbolic mutual information (wSMI), then deteriorated into the pathological range after the audiobook, without mood improvement. N1 showed mixed behavioral effects, with mood and quality of life improving after music. N3 showed no significant spatial improvement and worsened mood after music. These heterogeneous responses motivate larger studies of connectional predictors. The findings support the feasibility of classical music as an adjunct to neglect rehabilitation and suggest that cognitive changes may occur independently of mood, while the audiobook cannot be considered an inert control.

## Introduction

Spatial neglect is among the most disabling consequences of right-hemisphere stroke. Patients fail to detect, orient toward, or respond to stimuli in contralesional space (Bartolomeo, 2014), behaving as if the left part of the world did not exist. Left neglect occurs acutely in up to 61% of right-hemisphere strokes (Cazzoli et al., 2025), and about 40% of patients develop chronic neglect persisting for months (Overman et al., 2024); it predicts poor functional outcome, limited independence, and reduced quality of life (Chen et al., 2015). Converging evidence from lesion-symptom mapping, diffusion tractography, and resting-state imaging establishes neglect as a disorder of large-scale brain networks rather than of focal cortical dysfunction (Bartolomeo et al., 2007; Corbetta & Shulman, 2011; Doricchi et al., 2008; Toba et al., 2017, 2020, 2022). Damage to long-range white-matter tracts, especially the second and third branches of the superior longitudinal fasciculus (SLF II/III), predicts chronic neglect more reliably than cortical damage (Thiebaut De Schotten et al., 2014), and the relative integrity of these pathways, particularly the right SLF II and the inferior fronto-occipital fasciculus (IFOF), supports recovery (Kaufmann et al., 2024). Interhemispheric connectivity is equally critical: splenial disconnection predicts persistence, whereas preserved callosal connectivity is associated with spontaneous recovery and with response to prism adaptation (Bartolomeo, 2019; Kaufmann et al., 2024; Lunven et al., 2015, 2023; Lunven, Rode, Bourlon, Duret, Migliaccio, Chevrillon, Thiebaut De Schotten, et al., 2019).

The role of the intact left hemisphere in recovery has been debated, with proposals of either a maladaptive (Corbetta et al., 2005) or a compensatory (Bartolomeo, 2021; Bartolomeo & Thiebaut de Schotten, 2016) function. One way to reconcile these views is that the healthy hemisphere can compensate, but only when it can communicate with the lesioned hemisphere through preserved callosal pathways (Bartolomeo, 2019, 2021). This suggests that interventions promoting coordinated activity across the two hemispheres may aid neglect rehabilitation when the callosal pathways connecting them remain intact.

Intensive exposure to music is a possible means of promoting such communication. Music processing engages both hemispheres, with pitch and melody recruiting right temporal networks and rhythm recruiting left temporal and bilateral motor circuits (Peretz & Zatorre, 2005). Music-assisted rehabilitation improves cognitive recovery and mood after stroke (Särkämö et al., 2008) and induces connectivity changes after left-hemisphere stroke (Sihvonen et al., 2022). On this basis, Bartolomeo (2022) advanced three hypotheses: (H1) anatomo-functional connectivity is a major determinant of recovery, so chronic deficits should show signs of disconnection, and cognitive and connectivity improvements should co-vary; (H2) interhemispheric connectivity determines whether the healthy hemisphere is maladaptive or compensatory, with compensation requiring preserved callosal communication; and (H3) music can improve recovery by promoting intra- and interhemispheric communication, assessable through tractography and resting-state EEG biomarkers. Chea et al. (2024) first tested H3 in chronic post-stroke aphasia, comparing two weeks of daily classical music with standard care in four patients with left-hemisphere damage. In that pilot study, neuropsychological improvement was accompanied by increased theta-band weighted symbolic mutual information (wSMI), an EEG connectivity marker originally developed to index consciousness in non-communicative patients (King et al., 2013), and by increased fractional anisotropy, a quantitative marker of white-matter integrity, in callosal and intrahemispheric tracts. Whether classical music can likewise modulate connectivity and behavior after right-hemisphere stroke, in chronic neglect, remains unknown. Spatial neglect offers a particularly informative test case because evidence suggests that preserved callosal communication with the damaged right hemisphere may enable compensation by the intact left hemisphere (Bartolomeo, 2019, 2022; Lunven et al., 2015; Lunven, Rode, Bourlon, Duret, Migliaccio, Chevrillon, Thiebaut de Schotten, et al., 2019). Moreover, the weaker lateralization of attention compared with language (Bartolomeo & Seidel Malkinson, 2019) may leave greater compensatory potential in the contralesional hemisphere.

We report three patients with chronic left neglect, studied using Chea et al.’s (2024) protocol, with an audiobook as an active control for non-musical auditory stimulation and narrative engagement. We assessed white-matter microstructure (diffusion MRI tractography, principally of fronto-parietal, fronto-occipital, and callosal pathways) and functional connectivity (theta-band wSMI from resting-state high-density EEG), complemented by behavioral neglect tests and mood and quality-of-life measures to examine how potential changes in connectivity relate to changes in mood and well-being (Koelsch, 2014). Following H1–H3, we expected all patients to show signs of disconnection, behavioral responses to co-vary with connectivity changes, recovery to depend on callosal preservation, and theta wSMI to increase specifically after music in posterior attentional regions. We further expected individual differences to reflect each patient’s connectional profile rather than a uniform treatment effect.

## Methods

### Participants

Patients were recruited from the Rehabilitation Department of Pitié-Salpêtrière Hospital. Inclusion required a first-ever right-hemisphere stroke with persistent left neglect at least three months after onset. Exclusion criteria were impaired vigilance, confusion, global cognitive decline, severe psychiatric or prior neurological disease, and contraindications to MRI. All participants gave written informed consent. The study was promoted by Inserm (protocol C13-41) and approved by the IRB Île-de-France I.

Three right-handed men with ischemic strokes were enrolled (Table 1); none had prior musical training. N3 presented with clinically diagnosed chronic left spatial neglect, despite milder deficits than N1 and N2. N1 and N2 were neutral toward classical music, preferring jazz, rock, and rap, whereas N3 reported a strong pre-existing emotional and nostalgic attachment to it.

**Table 1.** Participant demographic and clinical characteristics. CBS, Catherine Bergego Scale; M, male.

| **Patient** | **N1** | **N2** | **N3** |
| --- | --- | --- | --- |
| **Age** | 63 | 48 | 64 |
| **Sex** | M | M | M |
| **Stroke onset (days)** | 155 | 173 | 196 |
| **Hospitalization status** | Inpatient rehabilitation | Outpatient rehabilitation | Outpatient rehabilitation |
| **Baseline CBS (max 30; higher scores indicating greater severity; pathological cut-off: 0)** | 15 | 13 | 2 |
| **Ongoing rehabilitation** | Physical and occupational therapy (daily)<br>Psychological support (weekly) | Physical and occupational therapy (daily)<br>Psychological support (weekly) | Physical and occupational therapy (3 times per week)<br>Psychological support (weekly) |
| **Study arm assignment** | A (music → audiobook) | A (music → audiobook) | B (audiobook → music) |

### Protocol

The crossover design used four assessment points (Figure 1). At inclusion (T0), a baseline neuropsychological assessment established severity; one week later (T1) it was repeated to confirm stability, and MRI and EEG were acquired. In Arm A (N1, N2), two weeks of music-assisted rehabilitation added to standard care preceded two weeks of audiobook listening added to standard care; in Arm B (N3), the order was reversed. Assessments were repeated after each block (T2, switch; T3, final). For N1 and N2, T2 followed music and T3 followed the audiobook; for N3, T2 followed the audiobook and T3 followed music. No procedural changes were made after the study began. No formal replication was planned within this three-case feasibility study; the present findings are intended to inform larger confirmatory studies. Treatment order was not randomized. No blinding was used, because participants and investigators could distinguish the audiovisual music and audiobook conditions.

**Figure 1.**
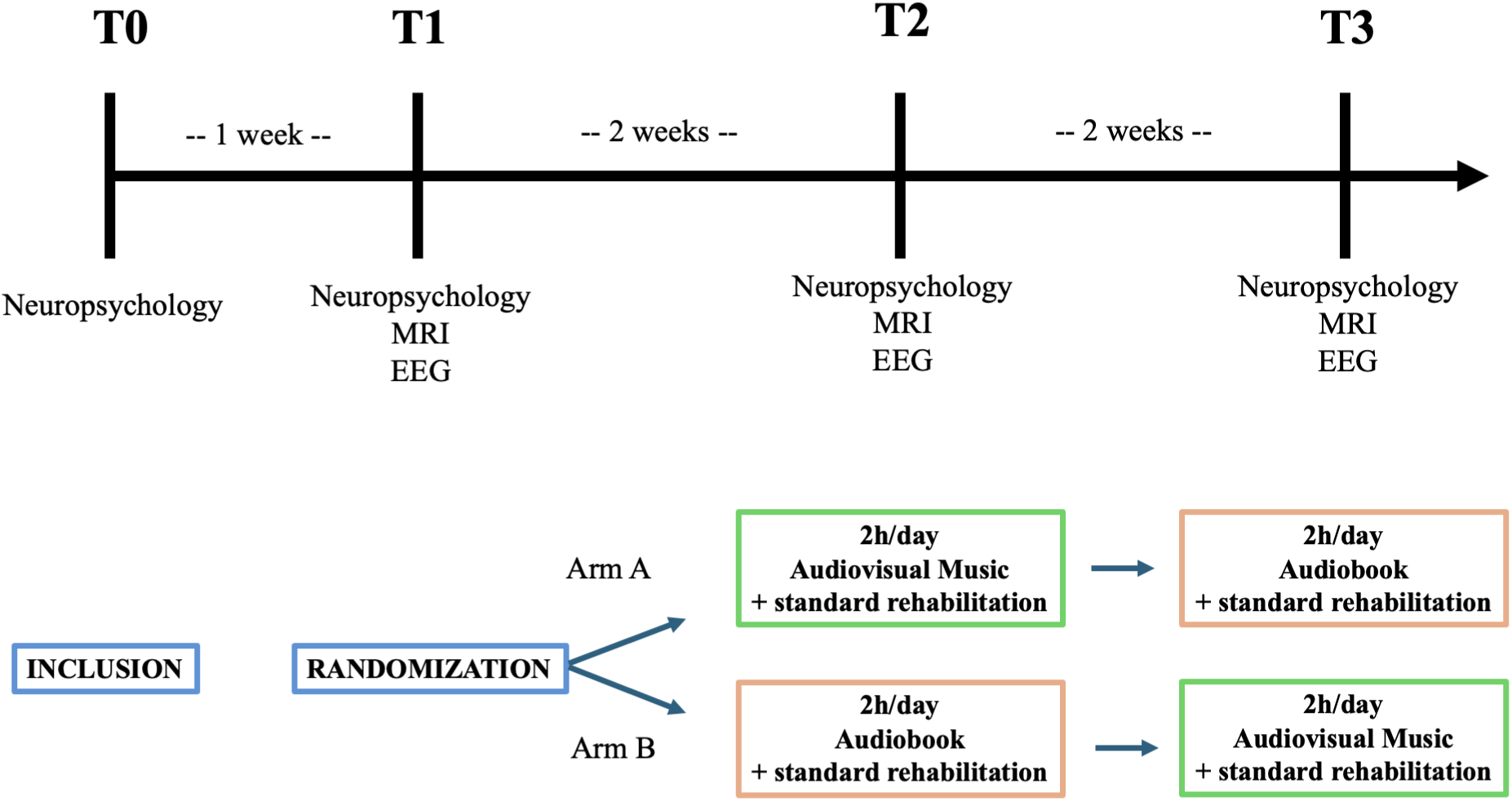
Crossover design: a multiple single-case approach with two intervention sequences (Arm A: music then audiobook; Arm B: audiobook then music) and four assessment time points (T0–T3). Each patient serves as their own control. Music and audiobook intervention

The music intervention, developed with Maestro Lorenzo Coppola and the Paris Mozart Orchestra, presented works by Haydn, Mozart, and Beethoven as audiovisual recordings, each preceded by a brief narrated historical and aesthetic introduction. The structure of Viennese classical music creates expectations over relatively extended phrases (typically 5–15 s), requiring perceptual representations to be actively maintained across time, a property argued to support sustained attention and bi-hemispheric processing in post-stroke rehabilitation (Bartolomeo, 2022). The same playlist was used for all patients, in 2-h sessions with a 15-min break after the first hour.

The audiobook (My Stroke of Insight, Jill Bolte Taylor; The Brain’s Way of Healing, Norman Doidge) served as an active auditory control, providing structured semantic content without musical structure. It was matched to the music condition in daily duration and total exposure (2 h/day on weekdays, 20 h over two weeks), but was delivered audio-only rather than audiovisually. All patients heard identical audiobook content in the same order.

### Neuropsychological, mood, and quality-of-life assessment

Neglect was assessed with the Neglect Assessment Battery (Batterie d’Évaluation de la Négligence, BEN; Azouvi et al., 2023), comprising Bells cancellation, figure copying, line bisection (5-cm and 20-cm tasks), clock drawing, reading, the overlapping-figures task, the Ogden figure copying task, and the CBS. We also administered the Mesulam (2000) and Albert (1973) cancellation tasks and an additional eight-line bisection task (Bartolomeo & Chokron, 1999; Toba et al., 2018). Mood was tracked with the French Profile of Mood States (POMS-f; Cayrou et al., 2003), from which Total Mood Disturbance (TMD) was derived (anxiety + depression + anger + fatigue + confusion − vigor). Quality of life was assessed with the French Stroke-Specific Quality of Life Scale (SS-QoL; Legris et al., 2018).

### Statistical analysis

Following Azouvi et al. (2023), three discriminative BEN subtests were analyzed as main outcomes: Bells cancellation, 20-cm line bisection, and reading. For the cancellation and reading tasks, each left-sided item was scored as a binary outcome (detected or omitted; read correctly or incorrectly). Within each patient, the outcome for each item was paired with the outcome for that same item at the next assessment (T1 vs T2 and T2 vs T3) and analyzed with the McNemar mid-p test (Fagerland et al., 2013), as in Chea et al. (2024); these paired comparisons included the 15 left-sided Bells items, the 30 left-sided items in each of the Mesulam and Albert tasks, and the 61 left-sided reading items. Effect sizes were Cohen’s d derived from probit-transformed paired proportions (Sánchez-Meca et al., 2003), with wide confidence bounds given the small item counts.

For 20-cm line bisection, deviations from the true midline were measured in millimeters across five lines, with positive values indicating rightward deviations and negative values indicating leftward deviations. Given the five observations per assessment, changes were analyzed descriptively in each patient by reporting mean deviation, the direction of change across individual lines, and crossings of the BEN cutoffs (+6.5 mm rightward; −7.5 mm leftward). POMS-f and SS-QoL were compared with the Brunner-Munzel test (Karch, 2021).

Inferential contrasts for cancellation, reading, POMS-f, and SS-QoL (T1 vs T2; T2 vs T3) were run separately per patient with Bonferroni correction. Results were interpreted jointly with effect sizes and BEN cut-off crossings. Line bisection and the remaining battery measures (8-line bisection, Ogden figure copying, clock drawing, overlapping figures) were interpreted descriptively.

### MRI acquisition and analysis

Structural and diffusion MRI used a Siemens 3.0 T Prisma Fit scanner (CENIR, Paris Brain Institute): T1-weighted MP2RAGE (1 mm isotropic) and multi-shell diffusion MRI with two phase-encoding directions, with 46 diffusion-weighted volumes at b = 1,500 s/mm² and 45 at b = 3,000 s/mm² per phase-encoding direction, plus seven non-diffusion volumes. Further acquisition details are provided in the Supplementary Materials. Lesion masks were drawn on the native T1 in MRIcron and normalized to MNI space with the SPM12 Clinical Toolbox (Rorden et al., 2012); gray-matter damage was quantified with the AAL3 atlas (Rolls et al., 2020). Diffusion data were preprocessed and fractional anisotropy (FA) maps were computed. Deterministic tractography was performed with manually delineated ROIs (Catani & Thiebaut De Schotten, 2012) for the SLF II, SLF III, IFOF, and corpus callosum, the tracts previously associated with neglect chronicity and recovery (Bartolomeo, 2019; Kaufmann et al., 2024; Lunven et al., 2015, 2023, 2019; Thiebaut De Schotten et al., 2014).

### EEG acquisition and analysis

Resting-state EEG was recorded at T1, T2, and T3 with a 256-channel EGI HydroCel HD net (250 Hz; 10 min, eyes closed). After removal of neck and face channels, 183 channels were analyzed. Following previous research (Chea et al., 2024), theta-band wSMI, a non-linear, information-theoretic functional-connectivity metric sensitive to states of consciousness and to brain damage (King et al., 2013), was computed with the NICE toolbox (Engemann et al., 2018) after preprocessing.

To compare the music and audiobook conditions across arms, wSMI was z-standardized channel-wise relative to the immediately preceding assessment (Arm A: music vs. baseline, audiobook vs. post-music; Arm B: audiobook vs. baseline, music vs. post-audiobook). Channel-level changes were tested with cluster-based permutation tests using a cluster-forming threshold of p < 0.05; significant clusters were visualized at p < 0.0001. Channel-to-channel edges were extracted from significant clusters only when the sign of change for an edge matched the sign of its cluster, and were rendered with Vizaj (Rolland & De Vico Fallani, 2023). Full analysis pipelines are described in the Supplementary Materials.

## Results

### Lesion mapping

Percentages of regional damage are summarized in Table S1 (Supplementary Materials), and lesion overlap is shown in Figure 2. N1 had a large right-hemisphere lesion (115.8 cm³) involving frontal, temporal, parietal, insular, and subcortical structures. N2 had a smaller right fronto-temporo-insular lesion (41.6 cm³). N3 had the smallest lesion (14.4 cm³), predominantly right temporo-parietal, with relative sparing of frontal and subcortical structures.

**Figure 2.**
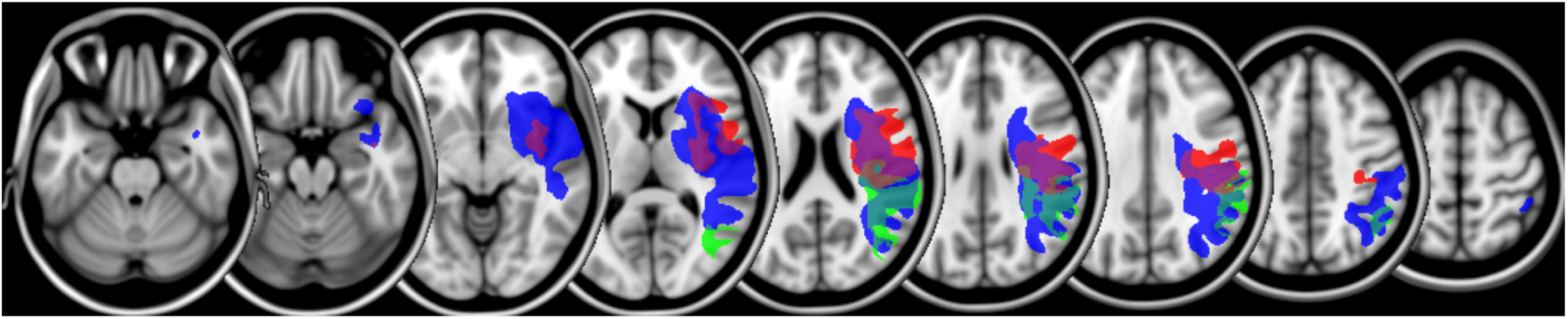
Baseline T1-weighted MRI lesion masks for N1 (blue), N2 (red), and N3 (green), shown on axial slices.

### White matter tractography

The right SLF II could not be reconstructed in any patient, consistent with it being the branch most consistently disrupted in chronic neglect (Thiebaut De Schotten et al., 2014) (Figure 3; Table S2, Supplementary Materials). The right SLF III and right IFOF were reconstructable only in N3, and the right SLF III showed partial disconnection. Left-hemisphere tracts were all reconstructable in all patients with comparable FA. The corpus callosum was preserved in all three patients, within the ranges reported in patients who responded to neglect rehabilitation (Lunven et al., 2015, 2019), with no evidence of splenial disconnection. Due to excessive head motion, only AP phase-encoded data were usable for N1.

**Figure 3.**
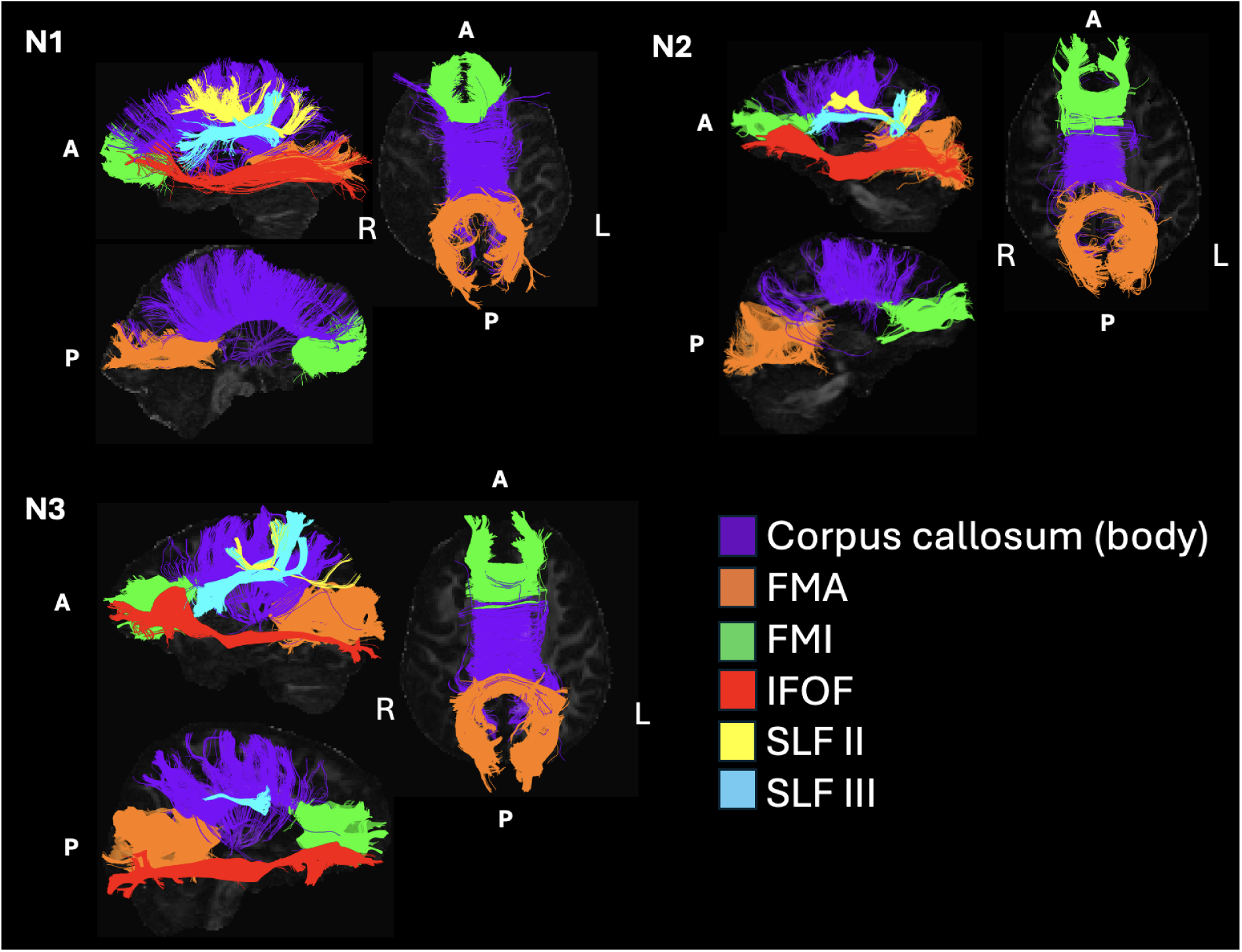
White-matter bundle reconstruction from ROI-based deterministic tractography in N1, N2, and N3. For each patient: left sagittal view showing the left superior longitudinal fasciculus (SLF) II (yellow), left SLF III (cyan), left inferior fronto-occipital fasciculus (IFOF, red), corpus callosum body (purple), forceps major (FMA, orange), and forceps minor (FMI, green); right sagittal view showing reconstructable right-hemisphere tracts of interest (none in N1 and N2; right SLF III and right IFOF in N3); and axial view of callosal sub-bundles. A = anterior; P = posterior; R = right; L = left.

### Neuropsychological outcomes

No adverse events occurred in any participant during either the music or audiobook intervention phase. BEN performance across the primary assessments is shown in Figure 4 and Table S3 (Supplementary Materials). Unless stated otherwise, cancellation and reading scores are reported as numbers of left-sided omissions, so that lower values indicate better performance.

**Figure 4.**
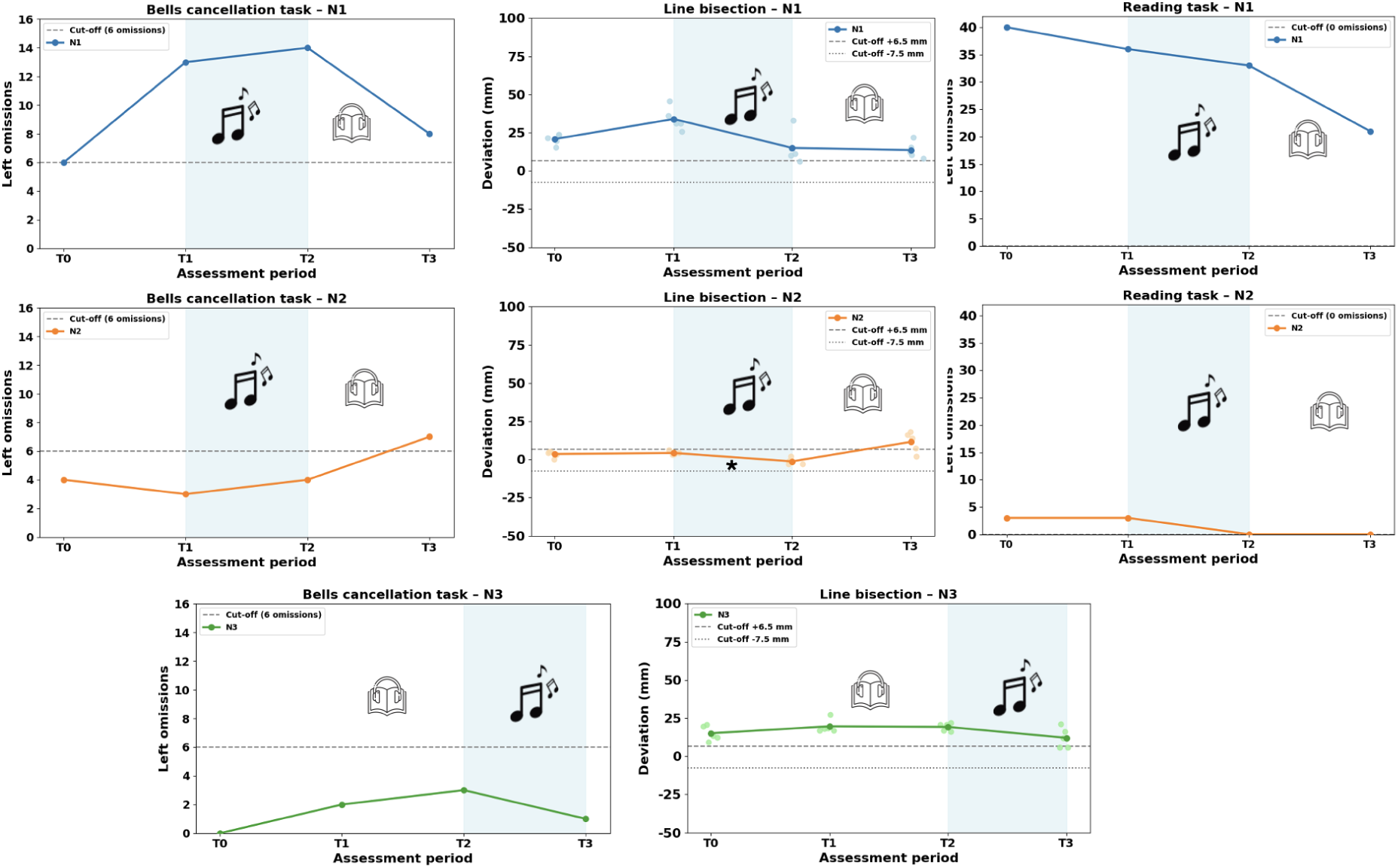
Neuropsychological performance across assessments for the three patients (N1–N3): left-sided omissions on Bells cancellation, line-bisection deviation (mm), and left-sided omissions on reading. Shaded regions indicate the music block. Statistical significance (* p_adj < 0.05, Bonferroni) applies only to cancellation and reading contrasts; line bisection is descriptive. N3 made no reading omissions.

N1 (Arm A: music → audiobook). On Bells, left-sided omissions did not change after music, but decreased after the subsequent audiobook (14 → 8; p_adj = 0.016, d = −0.904), remaining above cut-off. On Mesulam, omissions increased significantly after music (9 → 21; p_adj =0.001, d = −0.75). On Albert, omissions decreased after music (20 → 13; p_adj = 0.008, d =−0.59) and rose again after the audiobook (13 → 22; p_adj = 0.004, d = 0.64), remaining above cut-off throughout. All cancellation scores remained pathological across the protocol.

Reading omissions decreased significantly after the audiobook (33 → 21; p_adj = 0.003, d =−0.44), remaining above cut-off. Rightward deviation on 20-cm line bisection decreased after music (34.0 → 15.0 mm) but remained pathological; on the descriptive eight-line task, rightward deviation instead increased across the protocol (+30.0%, +38.1%, +46.3%; 3, 1, and 1 line omissions, respectively).

N2 (Arm A: music → audiobook). Left-sided cancellation asymmetry did not change significantly after either condition. On Mesulam, however, the right hemifield normalized after music (Left/Right hits, max 30/30: 21/21 → 26/30), while left-sided performance remained pathological both after music (9 → 4 omissions) and after the audiobook. On 20-cm line bisection, mean deviation shifted leftward after music from +4.2 to −1.2 mm, with all five lines moving in the same direction; both values were within normal limits. After the audiobook, deviation shifted rightward to a pathological +11.5 mm. The eight-line task showed the same pattern (+6.5% → −8.8% → +12%), with performance pathological after the audiobook (T3). Left-sided reading performance normalized after music, although one right-sided omission persisted (Table S3). Ogden figure copying normalized after music (1 → 0 omissions; FigureS1, Supplementary Materials). Clock drawing (Figure S2, Supplementary Materials) and overlapping figures were normal throughout. N3 (Arm B: audiobook → music). No comparison reached significance, but cancellation performance was not uniformly normal. Bells remained within the non-pathological range at all three assessments (Left/Right hits (max 15/15): 13/14 → 12/14 → 14/14 targets detected). On Mesulam (Left/Right hits, max 30/30), left-sided performance was above the pathological cut-off at baseline (T1: 27/29), normalized after the audiobook (T2: 30/29), and again exceeded the cut-off after music (T3: 27/30). On Albert (Left/Right hits, max 30/30), performance was within normal limits at baseline and after the audiobook (T1: 30/30 → T2: 30/30) but crossed the cut-off after music (T3: 30/27). Deviation on 20-cm line bisection remained pathological throughout, although it decreased after music (19.7 → 19.3 → 12.1 mm); the eight-line task showed small, stable deviations (+8.9%, +8.1%, +7.6%); and reading was error-free. Figure copying, clock drawing, and overlapping figures were unimpaired.

Overall, N2 showed the most consistent music-related changes, with improvement in reading and a leftward shift on line bisection; N1’s clearest gains followed the post-music audiobook block; and N3 showed no statistically significant improvement, although 20-cm bisection deviation decreased after music and cancellation performance varied.

### EEG functional connectivity

Theta-band wSMI differences between music and audiobook are shown in Figure 5. N1 showed a single cluster of increased wSMI over left middle and posterior regions, reflecting primarily intra-hemispheric left temporo-parietal and occipital coupling. N2 showed two opposing clusters: increased wSMI in occipito-parietal regions involving interhemispheric coupling, and decreased wSMI in bilateral frontal and central regions. N3 exhibited diffuse, widespread wSMI decreases with no discernible network organization or correspondence to attentional network topography, precluding a clear functional interpretation.

**Figure 5.**
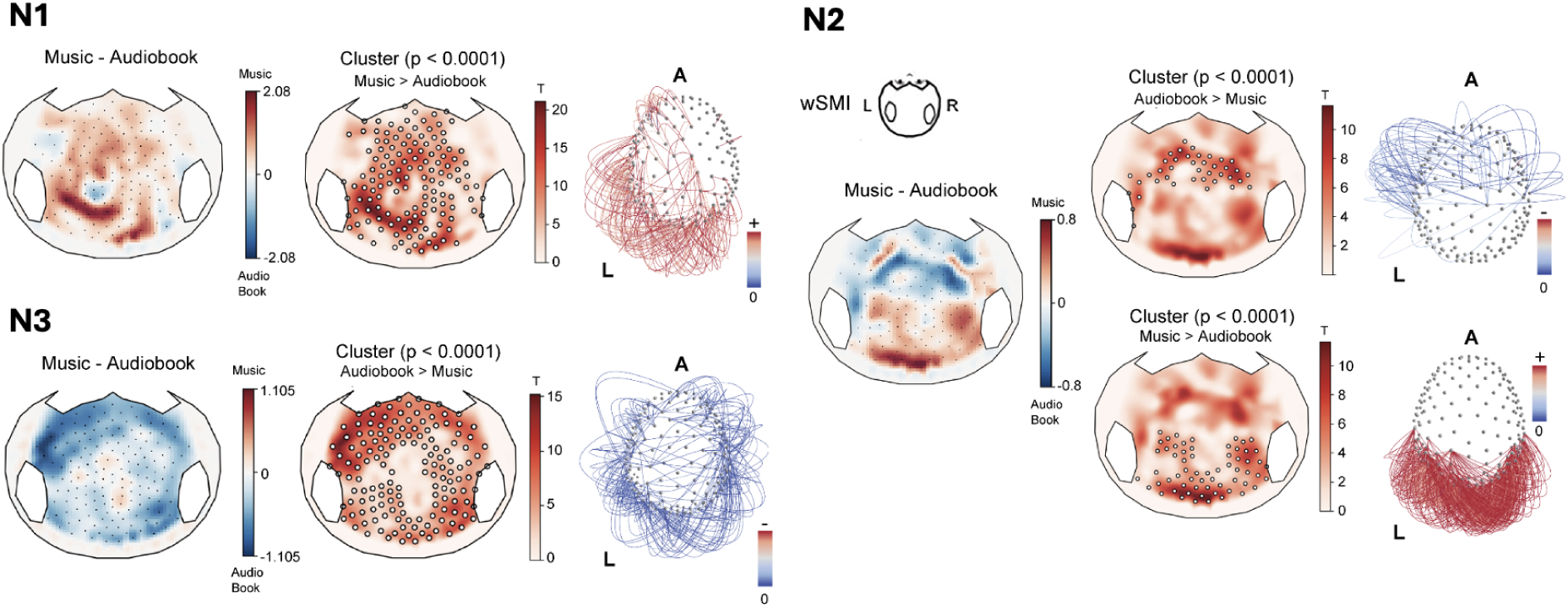
Theta-band weighted symbolic mutual information (wSMI) changes from pre- to post-music for N1, N2, and N3. For each patient: left, topographic map of significant channel clusters visualized at p < 0.0001; right, three-dimensional connectivity rendering of significant channel pairs, with edge color indicating t-value. For N2, both increased and decreased clusters are shown. wSMI changes were standardized to the mean and standard deviation of the immediately preceding assessment (see Methods). In the channel-to-channel renderings, the top 10% of edges are displayed for N1, which contained a large number of significant channels and channel pairs; for N2 and N3, all extracted edges are shown. Renderings were produced with Vizaj (Rolland & De Vico Fallani, 2023). L = left; R = right; A = anterior.

### Mood and quality of life

POMS-f TMD and SS-QoL total scores are shown in Figure 6; statistical results are given in Table S4 (Supplementary Materials). N1 improved significantly on both POMS-f (p_adj < 0.001) and SS-QoL (p_adj = 0.004) after music, with a significant SS-QoL regression after the audiobook (p_adj = 0.040). N2 showed no POMS-f change after either condition; SS-QoL was unchanged after music but improved after the audiobook (p_adj < 0.001). N3 showed significant mood deterioration after both conditions (p_adj = 0.029, p_adj = 0.026) and improved SS-QoL after music (p_adj = 0.024).

**Figure 6.**
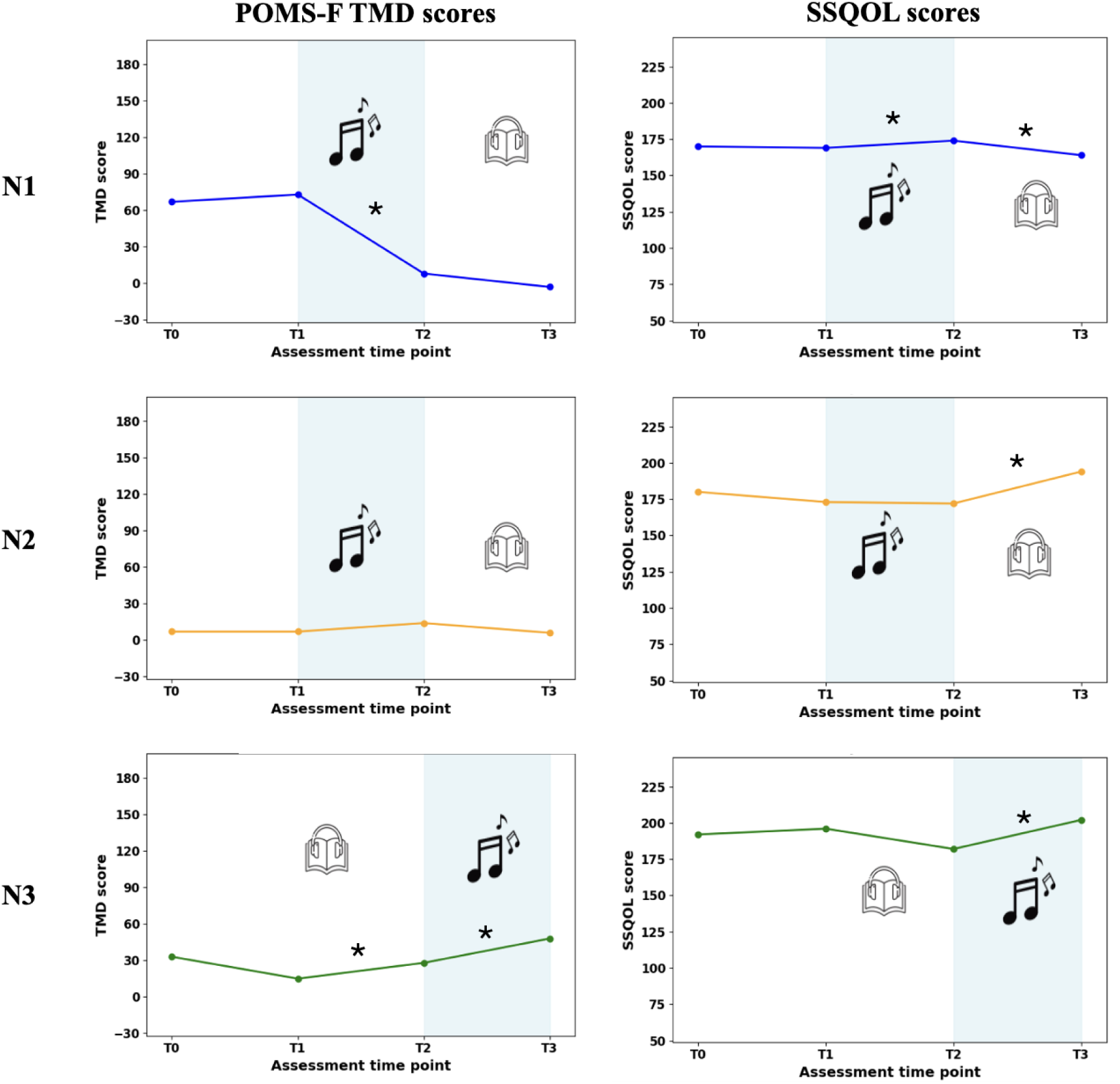
POMS-f Total Mood Disturbance (TMD) and Stroke-Specific Quality of Life (SS-QoL) total scores across assessments for N1 and N2 (Arm A) and N3 (Arm B). Shaded regions indicate the music block. Higher SS-QoL = better quality of life; higher POMS-f TMD = worse mood. * p_adj < 0.05 (Bonferroni).

Mood, quality-of-life, and spatial outcomes did not change uniformly. N1 showed mood and quality-of-life improvements after music, with spatial gains at different assessments; N2 improved on spatial measures after music without mood improvement; and N3 showed worsened mood, improved quality of life, and no statistically significant spatial improvement after music.

## Discussion

To our knowledge, this is the first multimodal study to examine whether sustained exposure to audiovisual classical music can modulate brain connectivity and behavior in chronic spatial neglect.

### Connectivity profiles as determinants of response (H1)

All three patients showed intrahemispheric disconnection in the damaged right hemisphere, consistent with the interpretation of spatial neglect as network-level dysfunction (Bartolomeo et al., 2007). The right SLF II could not be reconstructed in any patient, consistent with abundant previous evidence (Doricchi et al., 2008; Kaufmann et al., 2024; Lunven et al., 2015, 2019; Thiebaut De Schotten et al., 2014; Toba et al., 2022). Interhemispheric pathways, by contrast, were spared in every patient (see H2 below), so the disconnection observed here was specifically intrahemispheric.

Lesion volume and fronto-parietal disconnection nonetheless varied widely, from a large insular–parietal–basal ganglia lesion in N1 to a small temporo-parietal lesion with preserved right SLF III and IFOF in N3. This anatomical heterogeneity was paralleled by behavioral and EEG heterogeneity: N3, with the most preserved right fronto-parietal architecture, had the mildest deficit and the least room for improvement. With three cases, we cannot test the quantitative co-variation that H1 requires. The coexistence of anatomical and response heterogeneity motivates testing this relationship in larger samples (Lunven et al., 2019).

### Interhemispheric connectivity as a substrate for compensation (H2)

All three patients had a preserved corpus callosum (body, forceps major, and forceps minor), with FA in the ranges reported for responders to neglect rehabilitation (Lunven et al., 2015, 2019). This satisfies the structural prerequisite of the "competition to cooperation" framework, according to which left-hemisphere contributions to neglect compensation depend on preserved communication with the lesioned right hemisphere (Bartolomeo, 2019, 2021). Both patients who showed spatial changes also showed EEG changes involving posterior coupling, but with different topography and timing. N2’s gains followed music directly and were accompanied by increased posterior interhemispheric theta wSMI together with decreased fronto-central coupling. N1’s gains followed a mixed timeline, Mesulam worsening and Albert cancellation improving after music, but Bells and reading only after the subsequent audiobook, and his wSMI increase was confined to left posterior channels, a more lateralized pattern consistent with his larger lesion. This convergence supports H2, but it cannot establish that callosal integrity was necessary; that would require comparison with patients showing splenial or anterior callosal disconnection.

### Music as a modulator of connectivity (H3)

The strongest evidence comes from N2. His 20-cm line bisection shifted leftward across all five lines after two weeks of music (+4.2 to −1.2 mm), remaining within normal limits because baseline was already non-pathological, together with increased posterior interhemispheric theta wSMI. Bisection then deteriorated into the pathological range after the audiobook. This within-subject pattern parallels the observation of Chea et al. (2024) in aphasia, where music-related behavioral gains co-occurred with theta wSMI and white-matter changes. Convergence across two anatomically and cognitively distinct syndromes, left-hemisphere aphasia and right-hemisphere neglect, using the same intervention and the same EEG marker provides preliminary cross-syndrome support for music engaging a general connectivity-modulation mechanism.

#### Relation to prior music-based interventions in neglect

Soto et al. (2009) showed that preferred music improved attentional performance in three chronic neglect patients relative to unpreferred music or silence, associated in one patient with orbitofrontal–parietal fMRI coupling, and interpreted this as positive affect releasing attentional resources. Kaufmann et al. (2022) showed that preferred music combined with auditory spatial cueing reduced neglect more than music alone, with response predicted by the integrity of the right inferior parietal lobule, SLF II, and posterior callosal fibers. Our approach differs in three respects.

First, we used a fixed classical repertoire rather than preferred music, motivated by the structural properties of Viennese classical music (Haydn, Mozart, Beethoven), which balances predictable against surprising events over extended time spans and thereby places sustained demands on attention and working memory (Bartolomeo, 2022). Notably, N1 and N2, who expressed no particular affinity for classical music, nonetheless showed spatial change, whereas N3, who reported a strong nostalgic attachment to the repertoire, showed no statistically significant spatial improvement and declined in mood. These observations suggest that spatial change need not be accompanied by improved mood. N3’s pre-existing attachment to the repertoire does not establish that it induced positive affect during the intervention.

Second, our cumulative two-week exposure with pre/post assessment targets consolidated network change rather than the within-session, affect-driven modulation studied by Soto et al.; the regression seen in N2 raises the possibility that any benefit may require continued exposure, but does not establish a plasticity mechanism.

Third, resting-state theta wSMI indexes intrinsic coupling outside the task, complementing the task-related BOLD coupling reported by Soto et al. (2009). The two accounts need not compete: affective gating during listening and cumulative connectivity change may be complementary mechanisms (present results; Chea et al., 2024; Sihvonen et al., 2022).

#### Case-by-case interpretation

N2 showed the most coherent pattern. Line bisection shifted leftward after music on both the five-line and eight-line tasks, in parallel with increased posterior interhemispheric theta wSMI in regions implicated in spatial attention (Bartolomeo & Seidel Malkinson, 2019; Corbetta & Shulman, 2011), then reversed to the pathological range after the audiobook. Mesulam cancellation followed the same trajectory, with right-hemifield performance moving from pathological at T1 to normal after music and back to pathological after the audiobook. The concurrent fronto-central wSMI decrease may reflect reduced maladaptive frontal coupling (Takamura et al., 2026), or simply a lower cognitive-control demand associated with the behavioral shift; we treat it as hypothesis-generating. N2 showed no improvement in mood or quality of life after music, although quality of life improved after the audiobook. His spatial changes after music were therefore not accompanied by improvement on these measures. N1 complicates a straightforward reading of H3. His gains followed a mixed timeline: Mesulam worsened and Albert cancellation improved directly after music, whereas his clearest Bells and reading gains emerged only after the subsequent audiobook block. Several non-exclusive explanations are possible: delayed effects of music, consistent with increased left-posterior theta wSMI, or active engagement with the audiobook’s narrative content. Against a simple spatial interpretation, rightward deviation on the eight-line bisection task increased across the protocol (+15.2, +19.3, +23.4 mm), diverging from the 20-cm task. Spontaneous recovery is unlikely given the chronicity of neglect. These findings indicate that the audiobook cannot be treated as an inert control. N1’s mood and quality of life improved after music, suggesting that affective and network-level changes may coexist without producing uniform spatial benefit. N3 had the smallest lesion, the mildest clinically rated deficit, and preserved right SLF III and IFOF. No comparison reached statistical significance. Nevertheless, 20-cm bisection deviation decreased after music (19.3 to 12.1 mm), while remaining pathological. This numerical change suggests a possible task-specific improvement that was not statistically established. His eight-line bisection deviations were small and stable, and reading remained error-free. His diffuse EEG decreases have no clear functional interpretation; the worsening in mood may relate to his reported nostalgic response to the music.

### Limitations

The small sample and the heterogeneity in lesion profile, deficit severity, and arm assignment preclude definitive conclusions about efficacy. The present findings should therefore be regarded as preliminary, serving primarily to establish feasibility and to generate hypotheses. Both patients showing spatial changes (N1, N2) were in Arm A (music first), so treatment and order effects cannot be fully separated, although N2’s leftward shift after music followed by rightward regression after the audiobook argues against pure spontaneous recovery. Several factors also challenge the assumption that the audiobook is an inert control: N1’s clearest improvements occurred during the audiobook phase; the emotionally engaging texts may themselves recruit attentional and motivational processes; and the two conditions differed in sensory modality (audiovisual for music versus audio-only for the audiobook). We did not compare classical with preferred music (Soto et al., 2009), and so cannot test how affect- and connectivity-based mechanisms interact. The structural and EEG findings are correlational and cannot establish mediation. Standard tests may lack sensitivity when performance is already near ceiling, as on several tasks in N3, and eye-tracking (Kaufmann et al., 2020) or ecological measures (Cerrato et al., 2021; Kaufmann et al., 2023, 2025) would be better suited. Finally, all three patients had preserved callosal pathways, leaving open whether splenial or anterior callosal disconnection limits the response, as documented for prism adaptation (Lunven et al., 2015, 2019).

### Perspectives

Music may influence cognition through two partly dissociable routes, an affect-driven route involving orbitofrontal–parietal coupling during listening (Soto et al., 2009), and a connectivity-based route operating on cumulative timescales via posterior interhemispheric networks (present results; Chea et al., 2024), with their relative contribution depending on the patient, the music, and the temporal structure of the intervention.

Future music rehabilitation studies should stratify patients by connectional profile and test whether response is predicted by callosal and right fronto-parietal integrity, as reported for prism adaptation (Lunven et al., 2019); use factorial designs crossing classical with preferred music and cumulative with single-session exposure; and develop theta-band wSMI as a cost-effective, bedside-compatible biomarker of plasticity during rehabilitation.

## Conclusion

These three case studies provide initial multimodal evidence that two weeks of daily Viennese classical music can be feasibly delivered in chronic spatial neglect, and can be followed by changes in behavior and in theta-band resting-state functional connectivity. The clearest converging result, in N2, was a leftward shift across all five 20-cm lines after music, with a shift in the same direction on the descriptive eight-line task and subsequent deterioration into the pathological range after the audiobook, together with increased posterior interhemispheric theta wSMI. This parallels Chea et al. (2024) in aphasia and supports the prediction that music engages a connectivity-based mechanism relevant to post-stroke cognitive deficits in either hemisphere (Bartolomeo, 2022). The findings are compatible with a role for anatomo-functional connectivity in recovery and with the proposed importance of preserved callosal communication for compensation by the healthy hemisphere (Bartolomeo, 2019, 2021). The differing patterns of affective and cognitive change suggest that spatial improvement need not be accompanied by better mood, while affect-driven (Soto et al., 2009) and connectivity-based mechanisms may contribute jointly to recovery. Larger studies with connectional stratification, direct comparison of classical and preferred music, and combined online and cumulative outcome measures will be needed to establish efficacy. Until then, classical music listening remains a low-cost, well-tolerated, and theoretically grounded candidate adjunct in the treatment of chronic spatial neglect.

## Credit authorship contribution statement

RZ: Investigation, Formal analysis, Data curation, Software, Visualization, Writing – Original Draft, Writing – Review & Editing. YT: Methodology, Writing – original draft, Writing – review & editing, Visualization, Software, Validation, Formal analysis, Data curation. MC: Methodology, Writing – review & editing. LN: Methodology, Writing – review & editing. EB: Investigation, Supervision, Writing – review & editing. MT: Writing – review & editing, Validation, Supervision, Conceptualization. PB: Writing – review & editing, Validation, Supervision, Resources, Conceptualization.

## Acknowledgements

We thank Dr. François Stefanescu, Dr. Quentin Marcillière, Dr. Anna Jolu, and the Rehabilitation department team for their hospitality and support; Bintou Coulibaly, Sandra Coelho, and Camille Robert for their assistance with EEG acquisition at the Neurology Institute; and the CENIR platform for MRI acquisition. We also thank the patients and their families. R.Z. was the recipient of a doctoral fellowship from the Institute for Health Engineering (IUIS), Sorbonne University. Y.T. is supported by the Uehara Memorial Foundation Overseas Postdoctoral Fellowship and JSPS KAKENHI Grants (JP24K20519 and JP24KK0296). M.N.T’s work is supported by the ANR PRCI BRAINGAME. Research was promoted by Inserm (protocol C13-41), approved by the Ethical Committee Île-de-France I, and funded by the “Investissements d’avenir” programme ANR-10-IAIHU-06 and by the Fondation pour la Recherche Médicale (grant FR-AVC-017) to P.B.

## Conflict of interest

None

## Declaration of generative AI in the writing process

During the preparation and revision of this work, the authors used ChatGPT (OpenAI) and Claude (Anthropic) to assist with language editing, clarity, and manuscript consistency checks. The authors reviewed and edited the output and take full responsibility for the content of the publication.

## Notes

### Competing Interest Statement

The authors have declared no competing interest.

