## Supplementary materials for "Classical music in cognitive rehabilitation: Three case studies of spatial neglect"

Supplementary material

### Methods

### MRI acquisition and analysis

Structural and diffusion MRI data were acquired on a Siemens 3.0 T Prisma Fit scanner at the CENIR imaging center, Paris Brain Institute. T1-weighted images were obtained using an MP2RAGE sequence (TR = 5000 ms, TE = 3.24 ms, TI1/TI2 = 700/2500 ms, flip angles = 4°/5°, 1 mm isotropic voxels, matrix = 256 × 256, FOV = 256 mm, GRAPPA acceleration factor R = 3, 64-channel head-neck coil). Diffusion-weighted MRI used two phase-encoding directions and a multi-shell scheme, with seven non-diffusion volumes, 46 diffusion-weighted volumes at b = 1500 s/mm², and 45 at b = 3000 s/mm² per phase-encoding direction.
Lesion masks were drawn on native T1-weighted images using MRIcron. Native T1 images and lesion masks were normalized to MNI space using the SPM12 Clinical Toolbox (Rorden et al., 2012) with the enantiomorphic method (Nachev et al., 2008). Normalized masks were filtered to exclude voxels outside the brain or within the cerebrospinal fluid. Affected gray matter regions were quantified using the Automated Anatomical Labelling atlas version 3 (AAL3; Rolls et al., 2020).
Diffusion data were preprocessed in FSL (Jenkinson et al., 2012) using BET for brain extraction and TOPUP/EDDY for distortion, motion, and eddy-current correction. For one patient (N1), only AP phase-encoded data were usable owing to excessive head motion; this is acknowledged in the Results. Fractional anisotropy (FA) maps were computed using StarTrack (Dell'Acqua et al., 2013). Deterministic tractography was performed using

StarTrack with FA termination threshold = 0.2, angle threshold = 35°, and step size = 0.5 mm. Tracts of interest were extracted in TrackVis (Wang et al., 2007) using a region-of-interest approach, with ROIs manually delineated according to the anatomical guidelines of Catani and Thiebaut de Schotten (2008; 2012). The following tracts were dissected bilaterally where reconstructable: the second and third branches of the superior longitudinal fasciculus (SLF II, SLF III), the inferior fronto-occipital fasciculus (IFOF), and the corpus callosum body, forceps major, and forceps minor.

### EEG acquisition and analysis

Resting-state EEG was recorded at T1, T2, and T3 in the Department of Neurology of Pitié-Salpêtrière Hospital using a 256-channel EGI HydroCel HD net (sampling rate 250 Hz, electrode impedance < 45 kΩ). Recordings lasted 10 minutes with eyes closed. Patients were monitored throughout for drowsiness or eye opening.

EEG preprocessing was performed in Python using MNE-Python (Gramfort et al., 2013), the NICE tools (Engemann et al., 2018), and custom scripts. Continuous data were band-pass filtered (0.5 Hz, 6th-order Butterworth; 45 Hz, 8th-order Butterworth) and segmented into 1-s epochs (−200 to 800 ms) with a 550–850 ms random jitter, matching the wSMI protocol (Bourdillon et al., 2020; King et al., 2013). The segmentation time was set to the start of the recording.

Automated artifact rejection was used for both amplitude- and statistics-based criteria. Channels with peak-to-peak amplitude exceeding 100 µV in more than 50% of epochs were marked as bad. Channels with z-score > 4 on variance were iteratively rejected (up to four iterations). Epochs were rejected if more than 10% of remaining channels exceeded the 100 µV threshold. A temporary copy of the data was then high-pass filtered at 25 Hz (4th-order zero-phase Butterworth), and a further iterative z-score procedure was applied to the high-frequency standard deviation to identify high-frequency artifacts. The resulting rejection criteria were applied to the original broadband data used for subsequent analyses. Quality control required retention of at least 30% of epochs and 70% of channels. Data were re-referenced to the common average. Bad channels were interpolated using spherical spline interpolation (Perrin et al., 1989). Baseline correction was applied using the −200 to 0 ms interval.

Theta-band weighted symbolic mutual information (wSMI) was computed as the connectivity marker of interest, following Chea et al. (2024). wSMI is a non-linear functional connectivity metric grounded in information theory that has shown sensitivity to states of consciousness and to brain damage (Birba et al., 2021; Engemann et al., 2018; King et al., 2013; Sitt et al., 2014). EEG signals were low-pass filtered at 10.42 Hz (corresponding to $\tau = 8$ [32 ms], $m = 3$) and current source density estimation was applied via pyCSD (https://github.com/nice-tools/pycsd) to attenuate volume conduction. Channel-wise wSMI features were obtained by taking the median across retained epochs. The 73 outer-perimeter electrodes covering non-scalp facial regions were excluded as nuisance channels, leaving 183 channels for analysis.

To allow comparison of music and audiobook effects across the two study arms, in which the same condition is preceded by different reference time points, wSMI values were z-standardized channel-wise to the mean and standard deviation of the immediately preceding assessment.

For patients N1 and N2, assigned to Arm A, standardized wSMI changes were computed as (Music−Baseline mean)/Baseline SD and (Audiobook−Music mean)/Music SD. For Patient N3, assigned to Arm B, standardized wSMI changes were computed as (Audiobook−Baseline mean)/Baseline SD and (Music−Audiobook mean)/Audiobook SD. The reference mean and standard deviation were computed across all retained epochs of the preceding session. The z-scored channel wSMI then constituted the observations for the statistical comparison.

Channel-level changes between the music and audiobook conditions were assessed using cluster-based permutation tests following Maris & Oostenveld (2007) implemented in

MNE-Python (Gramfort et al., 2013). We used 10,000 permutations of the condition labels, two-sample Welch's t-tests, and a cluster-forming threshold of $p < 0.05$. Significant clusters were visualized using a more stringent threshold of $p < 0.0001$, following Chea et al. (2024). To visualize channel-to-channel connectivity, we extracted edges from significant clusters only when the direction of change of each edge was consistent with the sign of the corresponding significant cluster. These edges were visualized using Vizaj (Rolland & De Vico Fallani, 2023)

### Results

***Supplementary materials Table* S1**. Percentage of regional lesion damage based on the AAL3 atlas (Rolls et al., 2020). The AAL3 (Automated Anatomical Labelling, version 3) is an atlas that parcellates the cortex and subcortical structures into anatomically defined regions, allowing the extent of lesion damage to be quantified within each region.

| **Brain Region (Right hemisphere)** | **N1** | **N2** | **N3** |
|---|---|---|---|
| Precentral Gyrus | 0 | 1.42 | 0 |
| Middle Frontal Gyrus | 0.01 | 0 | 0 |
| Inferior Frontal Gyrus, pars operculis | 11.28 | 17.00 | 0 |

| | | | |
|---|---|---|---|
| Right Inferior Frontal Gyrus, pars triangulis | 3.68 | 2.74 | 0 |
| Inferior Frontal Gyrus, pars orbitalis | 13.23 | 0 | 0 |
| Rolandic Operculum | 58.38 | 67.26 | 7.16 |
| Olfactory Cortex | 4.506 | 0 | 0 |
| Gyrus Rectus | 0.15 | 0 | 0 |
| Insula | 98.45 | 42.44 | 2.07 |
| Hippocampus | 0.70 | 0 | 0 |
| Amygdala | 23.87 | 0.51 | 0 |
| Superior Occipital Gyrus | 3.41 | 0 | 0 |
| Middle Occipital Gyrus | 13.20 | 0 | 8.36 |
| Postcentral Gyrus | 6.71 | 13.62 | 0 |
| Superior Parietal Gyrus | 0.58 | 0 | 0 |
| Inferior Parietal Gyrus | 34.53 | 0 | 9.29 |

| | | | |
|---|---|---|---|
| Supramarginal Gyrus | 56.06 | 17.31 | 27.12 |
| Angular Gyrus | 47.87 | 0 | 20.57 |
| Caudate Nucleus | 20.45 | 0 | 0 |
| Putamen | 93.23 | 37.39 | 0 |
| Globus Pallidus | 81.35 | 2.70 | 0 |
| Thalamus | 1.10 | 0 | 0 |
| Heschl's Gyrus | 93.39 | 26.19 | 0.26 |
| Superior Temporal Gyrus | 71.37 | 3.78 | 9.39 |
| Temporal Pole, Superior Temporal Gyrus | 19.57 | 0.04 | 0 |
| Middle Temporal Gyrus | 14.41 | 0 | 11.09 |

***Supplementary materials table* S2**. Baseline fractional anisotropy (FA) of the bilateral inferior fronto-occipital fasciculus (IFOF), bilateral branches II and III of the superior longitudinal fasciculus (SLF II/III), the body of the corpus callosum, forceps major (FMA), and forceps minor (FMI). N/A: not available; tract could not be reconstructed.

| Tract | N1 | N2 | N3 |
|---|---|---|---|
| IFOF left | 0.44 | 0.44 | 0.48 |
| IFOF right | N/A | N/A | 0.41 |
| SLF II left | 0.37 | 0.36 | 0.37 |
| SLF II right | N/A | N/A | N/A |
| SLF III left | 0.39 | 0.41 | 0.40 |
| SLF III right | N/A | N/A | 0.29 |

| | | | |
|---|---|---|---|
| Corpus Callosum (body) | 0.42 | 0.45 | 0.46 |
| FMA | 0.50 | 0.48 | 0.52 |
| FMI | 0.38 | 0.35 | 0.32 |

**Supplementary Table S3. Individual neuropsychological performance across the test battery at baseline and post-intervention timepoints for each patient. Scores are shown for each assessment period together with adjacent-timepoint comparisons (T1 vs T2, T2 vs T3). Left-sided omissions on the Bells, Mesulam, Albert, and text-reading tasks were compared using the McNemar mid-p test. Line bisection (20 cm) and all other measures (8-line bisection, Ogden figure copying, clock drawing, overlapping figures) are reported descriptively.**

P-values were Bonferroni-corrected across the two adjacent inferential comparisons within each patient. Effect sizes for cancellation and reading are reported as Cohen's d; a positive d represents more hits. No inferential p-values or effect sizes are reported for line bisection. Asterisks (*) denote scores meeting the pathological cut-off indicated for each test. Note that patient N3 received the audiobook condition before music (T2 audiobook, T3 music), the reverse order to N1 and N2. R: right; L: left; C: centre; N/A: effect size undefined (no discordant pairs). *p_adj < 0.05 (Bonferroni).* In the 8-line bisection task, left-lateralised omitted lines were calculated as 100% rightward deviation, right-lateralised omitted lines were not counted.

| Test (cut-off) | N1 | N2 | N3 |
|---|---|---|---|
| **Protocol / condition order** | T1 Baseline → T2 Music → T3 Audiobook | T1 Baseline → T2 Music → T3 Audiobook | T1 Baseline → T2 Audiobook → T3 Music |

| | | | |
|---|---|---|---|
| **Bells cancellation (L/R hits, max=15/15) Cut-off: >6 omissions on either hemifield; (L−R)>2; first column >5; time >183s** | T1 2/14*, 1st column 7*, 210s* T2 1/14*, 1st column 7*, 148s T3 7/15*, 1st column 7*, 235s* T1→T2: p=1.000, d=0.146 T2→T3: p=0.016, d=−0.904 | T1 12/12*, 1st column 7*, 87s T2 11/14*, 1st column 7*, 90s T3 8/13*, 1st column 7*, 60s T1→T2: p=1.000, d=0.146 T2→T3: p=0.578, d=0.295 | T1 13/14, 1st column 1, 180s T2 12/14, 1st column 1, 102s T3 14/14, 1st column 1, 150s T1→T2: p=1.000, d=0.112 T2→T3: p=0.500, d=−0.379 |
| **Mesulam cancellation (L/R hits, max=30/30) Cut-off: >0 omission per hemifield if < 50 yo; >1 if >50** ; >4 omissions per hemifield if >80 yo (Lowery et al., 2004; Mesulam, 2000) | T1: 21/23* (90s) T2: 9/29* (150s) T3: 14/30* (273s) T1→T2: p=0.001, d=−0.75 T2→T3: p=0.292, d=−0.281 | T1: 21/21* (81s) T2: 26/30* (85s) T3: 25/29* (81s) T1→T2: p=0.078, d=−0.413 T2→T3: p=0.578, d=0.208 | T1: 27/29* (109s) T2: 30/29 (104s) T3: 27/30* (95s) T1→T2: p=0.25, d=−0.327 T2→T3: p=0.25, d=0.327 |
| **Albert cancellation (L/R hits, max=30/30)** | T1: 10/29* (40s) T2: 17/29* (40s) T3: 8/30* (82s) | T1: 29/30* (33s) T2: 29/29* (42s) T3: 30/29* (40s) | T1 30/30 (41s) T2 30/30 (32s) T3 30/27* (30s) |

| | | | |
|---|---|---|---|
| **Cut-off: >0 omission & >70% on left** (Albert, 1973) | T1→T2: p=0.008, d=−0.592<br>T2→T3: p=0.004, d=0.643 | T1→T2: p=1.000, d=0<br>T2→T3: p=1.000, d=−0.18 | T1→T2: p=1.000, d=N/A<br>T2→T3: p=1.000, d=N/A |
| **Text reading (L/R hits, max=61/55) Cut-off: >0 omission; (L−R)≠0** (Azouvi et al., 2002) | T1: 25/47*<br>T2: 28/55*<br>T3: 40/55*<br>T1→T2: p=0.131, d=−0.248<br>T2→T3: p=0.003, d=−0.438 | T1: 58/54*<br>T2: 61/54*<br>T3: 61/54*<br>T1→T2: p=0.500, d=−0.182<br>T2→T3: p=1.000, d=N/A | T1: 61/55<br>T2: 61/55<br>T3: 61/55<br>T1→T2: p=1.000, d=N/A<br>T2→T3: p=1.000, d=N/A |
| **20-cm line bisection (mm deviation ± SD) Cut-off: >+6.5 mm or < −7.5 mm** (Azouvi et al., 2002) | T1: 34±7.5*<br>T2: 15±10.6*<br>T3: 13.7±5.25* | T1: 4.3±1.2<br>T2: −1.2±2.1<br>T3: 11.5±6.7* | T1: 19.7±4.4*<br>T2: 19.3±2.5*<br>T3: 12.1±6.7* |
| **8-lines bisection Cut-off: 11.1%** (Toba et al., 2018) | T1: +15.19 mm* / +30.0%<br>Omissions: 3<br>T2: +19.31 mm* / +38.1%<br>Omission: 1 | T1: +3.31 mm / +6.5%<br>Omission: 0<br>T2: −4.44 mm / −8.8%<br>Omission: 0 | T1: +4.50 mm / +8.9%<br>Omission: 0<br>T2: +4.13 mm / +8.1%<br>Omission: 0 |

| | | | |
|---|---|---|---|
| | T3: +23.43 mm* / +46.3% Omission: 1 | T3: +6.06 mm* / +12.0% Omission: 0 | T3: +3.88 mm / +7.6% Omission: 0 |
| **Ogden figure copying Cut-off: >0 omission; time >190s** (Azouvi et al., 2002) | T1: 2* (90s) T2: 2* (100s) T3: 2* (92s) | T1: 1* (60s) T2: 0 (50s) T3: 0 (65s) | T1: 0 (132s) T2: 0 (58s) T3: 0 (65s) |
| **Clock drawing Cut-off: >0 omission; time >70s** (Azouvi et al., 2002) | T1: 1* (30s) T2: 0 (40s) T3: 0 (33s) | T1: 0 (27s) T2: 0 (32s) T3: 0 (30s) | T1: 0 (22s) T2: 0 (15s) T3: 0 (20s) |
| **Overlapping figures (L/R hits, max=5/5 ; Position of the first hit item, Right/Center/Left) Cut-off: >0 omission (L−R)** (Azouvi et al., 2002) | T1: 4/5(R)*, 4/5(R*), 5/5(R), 3/5(R)*, 3/5(R) * T2: 3/5(R) *, 4/5(R) *, 3/5(R) *, 3/5(R) *, 4/5(R) * T3: 4/5(R) *, 4/5(R) *, 4/5(R) *, 3/5(R)*, 5/5(R) | T1: 5/5(C), 5/5(C), 5/5(C), 5/5(C), 5/5(C) T2: 5/5(C), 5/5(C), 5/5(C), 5/5(R), 5/5(C) T3: 5/5(L), 5/5(R), 5/5(C), 5/5(C), 5/5(C) | T1: 5/5(L), 5/5(C), 5/5(C), 5/5(C), 5/5(C) T2: 5/5(R), 5/5(R), 5/5(R), 5/5(C), 5/5(R) T3: 5/5(R), 5/5(R), 5/5(C), 5/5(R), 5/5(R) |

***Supplementary materials* table S4.** Brunner-Munzel statistical results on patients' subjective assessment of mood (POMS) and quality of life (SS-QoL) across assessment periods. Statistical tests were performed by pairing individual questionnaire items across assessment periods.

| *Patient* | *Assessment period* | *POMS* | *SS-QoL* |
|---|---|---|---|
| *N1* | *T1 vs T2* | p_adj < 0.001 | p_adj = 0.004 |
| | *T2 vs T3* | p_adj = 0.612 | p_adj = 0.040 |
| *N2* | *T1 vs T2* | p_adj = 0.113 | p_adj = 0.365 |
| | *T2 vs T3* | p_adj = 1.000 | p_adj < 0.001 |
| *N3* | *T1 vs T2* | p_adj = 0.029 | p_adj = 0.170 |
| | *T2 vs T3* | p_adj = 0.026 | p_adj = 0.024 |

***Figure S1. Patient N2's Ogden figure copying at each timepoint***

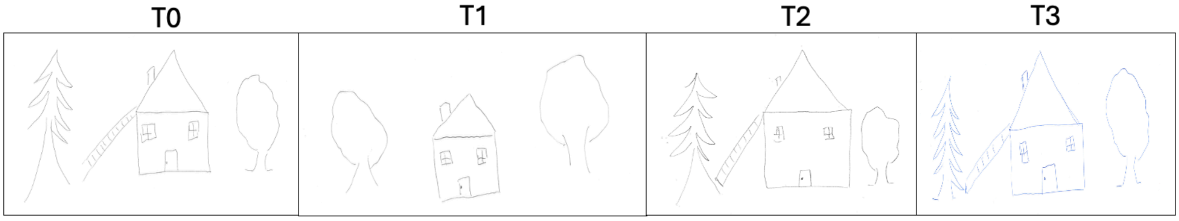


***Figure S2. Patient N2's clock drawing at each timepoint***

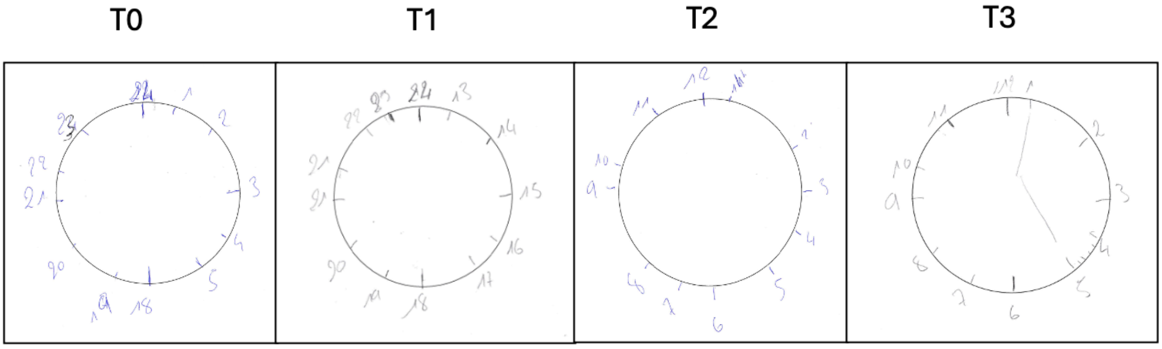
